# Stochastic model of BKPy Virus replication and assembly

**DOI:** 10.1101/746149

**Authors:** Suzy M. Stiegelmeyer, Liesl K. Jeffers-Francis, Morgan C. Giddings, Jennifer Webster-Cyriaque

## Abstract

BK Polyomavirus (BKPyV), belongs to the same family as SV40 and JC Virus and has recently been associated with both Sjögrens Syndrome and HIV associated Salivary Gland Disease. BKPyV was previously only known for causing the rejection of kidney transplants. As such, BKPyV infection of salivary gland cells implicates oral transmission of the virus. BKPyV replicates slowly in salivary gland cells, producing infectious virus after 72-96 hours. However, it remains unclear how this virus infects or replicates within salivary gland cells, blocking the development of therapeutic strategies to inhibit the virus. Thus, an intracellular, computational model using agent-based modeling was developed to model BKPyV replication within a salivary gland cell. In addition to viral proteins, we modeled host cell machinery that aids transcription, translation and replication of BKPyV. The model has separate cytosolic and nuclear compartments, and represents all large molecules such as proteins, RNAs, and DNA as individual computer “agents” that move and interact within the simulated salivary gland cell environment. An application of the Boids algorithm was implemented to simulate molecular binding and formation of BKPyV virions and BKPyV virus-like particles (VLPs). This approach enables the direct study of spatially complex processes such as BKPyV virus self-assembly, transcription, and translation. This model reinforces experimental results implicating the processes that result in the slow accumulation of viral proteins. It revealed that the slow BKPyV replication rate in salivary gland cells might be explained by capsid subunit accumulation rates. BKPyV particles may only form after large concentrations of capsid subunits have accumulated. In addition, salivary gland specific transcription factors may enable early region transcription of BKPyV.

## Introduction

BKPyV, a polyomavirus family member, is a non-enveloped, small, double-stranded DNA virus. The BKPyV genome codes for structural (VP1, VP2, VP3) and non structural (agnoprotein, small and large t antigen) genes whose transcription is controlled by a regulatory region. BKPyV is believed to cause a harmless latent infection in healthy people but may reactivate if the immune system has been compromised (Padgett and Walker, 1976). BKPyV is known to cause BKPyV nephropathy (BKN) a kidney transplant complication where reactivated BKPyV induces cell necrosis due to immunosuppressive drug regimens (Nickeleit et al., 2003). BKPyV sequences have been found in many organs in the human body—kidneys, liver, stomach, lungs, parathyroid glands, lymph nodes, tonsils, lymphocytes, bladder, prostate, uterine cervix, vulva, lips and tongue (Tognon et al., 2003). Recently, BKPyV has been detected in HIV positive patients with HIV associated salivary gland disease (HIV SGD) and shown capable of reproducing in salivary gland cells (Burger-Calderon et al., 2014; Jeffers et al., 2009). Since salivary gland diseases such as HIV SGD or Sjögren’s Syndrome do not have a known etiological agent, the association with BKPyV is intriguing and thus the study of the BKPyV replication process can aid in the understanding of the pathogenesis of the disease. We are pursuing this relationship with BKPyV using a computational model to reproduce the replication and assembly of BKPyV within a salivary gland cell.

Much of what is understood regarding BKPyV cell entry and intracellular trafficking have been from Monkey kidney Vero cells. However, recently, studies from human renal proximal tubular epithelial cells (HRPTEC) have been conducted since tubular epithelial cells are the main natural target of BKPyV infection (Moriyama and Sorokin, 2008). Compared to Vero cells, the BKPyV course of infection in HRPTEC seems to be relatively slow with the process taking at least 24–48 h. Pastrana et al identified distinct BKPyV genotypes with different cellular tropisms suggesting a potential for a BKPyV genotype preference for salivary gland cells (Pastrana et al., 2013). While we have recently determined that salivary gland cells are permissive for BKPyV replication, BKPyV replication is even slower when compared to HRTECs, with the process taking at least 72 hours (Burger-Calderon et al., 2014; Jeffers et al., 2009).

In the kidney cell, BKPyV is believed to enter the cell through caveolae-mediated endocytosis (Eash et al., 2004; Moriyama et al., 2007) after binding with ganglioside GD1b or GT1b (Dugan et al., 2005; Low et al., 2006) on the cell surface. Recently, the same receptor was confirmed for entry into salivary gland cells (Jeffers et al., 2009). It was determined that BKPyV may enter an acidic compartment after entry (Jiang et al., 2009). It is then believed to use the cell’s cytoskeleton (Eash and Atwood, 2005; Jiang et al., 2009; Moriyama and Sorokin, 2008) where it is transported to the ER, bypassing the Golgi, eventually gathering in the perinuclear region (Drachenberg et al., 2003; Inoue et al., 2015). Gathering in the perinuclear region was also observed in salivary gland cells (Jeffers et al., 2009). In HRPTECs, BKPyV particles are found in the ER at 6–8 hours after infection (Moriyama and Sorokin, 2008). BKPyV disassembly occurs due to VP1 cleavages prior to reaching the ER (Jiang et al., 2009). The nuclear localization signal located on VP2 and VP3 capsid proteins aid BKPyV entry into the nucleus where viral replication occurs (Bennett et al., 2015). BKPyV egress may occur by cell lysis but BKPyV virions have also been observed in vesicles in the cytoplasm (Drachenberg et al., 2003). The agnoprotein is also believed to play a role in nuclear egress (Johannessen et al., 2008; Okada et al., 2005) including acting as a viroporin (Suzuki et al., 2010) and by possibly inhibiting cell proliferation when interacting with PCNA, a gene involved in DNA replication (Gerits et al., 2015).

We assume that the steps in the BKPyV life cycle within kidney cells are very similar to salivary gland cells. See Figure 1 for a representation of BKPyV entry into a salivary gland cell. There is much unknown about the BKPyV life cycle in salivary gland cells. Thus, we developed an intracellular model of BKPyV replication and assembly within a salivary gland cell to assess BKPyV assembly. Viral transcription and replication using host cell machinery is modeled leading to the eventual assembly of virus like particles (VLP) as well as BKPyV virions to capture viral rates of production. By reproducing the BKPyV replication process, we will be able to understand the viral replication process, the progression of disease and how to inhibit it.

**Figure 1.**
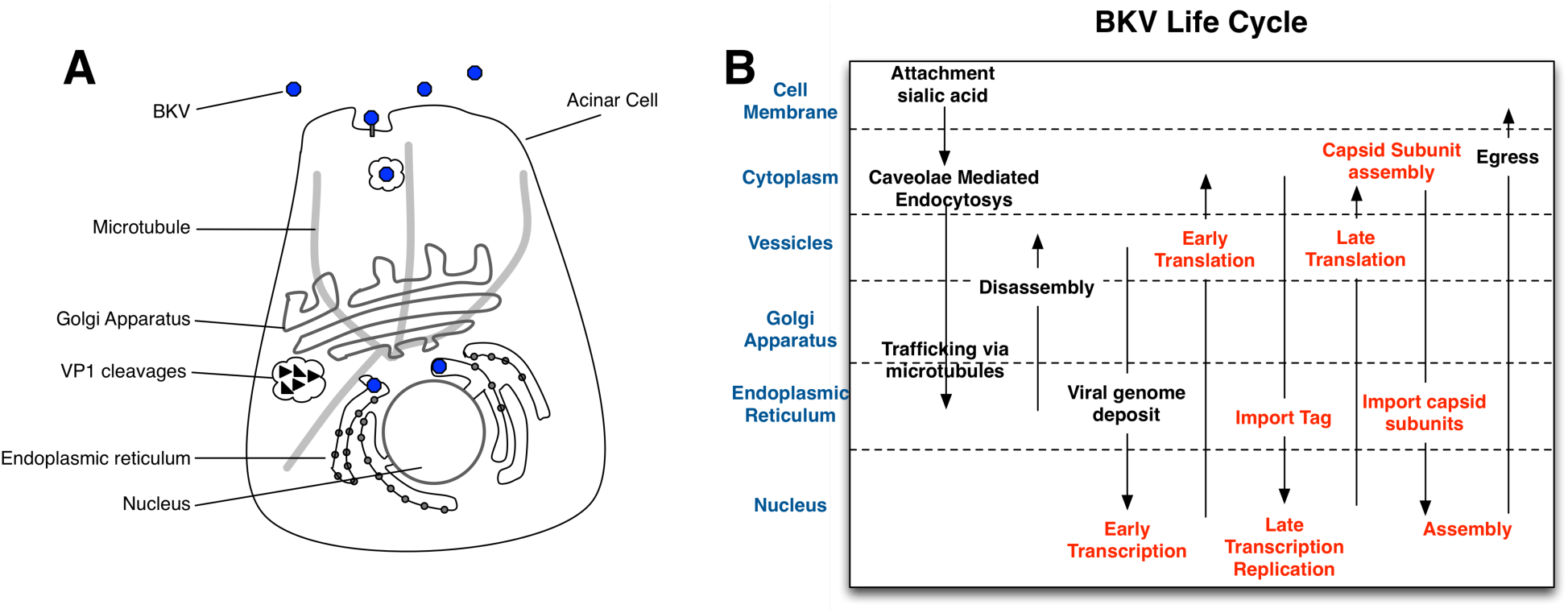
A) Mock-up of BKPyV entry into a salivary gland cell. B) Biological and computational model of the BKPyV Life Cycle reading from left to right. The portions of the viral lifecycle incorporated in the model are indicated in red. The arrows indicate transport of components from one compartment to the next.

Traditionally, the majority of viral pathogenesis models are mathematically based using differential equations. The models represent host cell and virus interaction, modeling only virus and host cell concentrations (Kepler et al., 2007; Perelson et al., 1996; Ribeiro et al., 2002). These models did not examine intracellular viral function or replication at the molecular level. Complex structures, patterns and phenotypes in biology are often the result of biochemical interactions between molecules. The T=7 icosahedral structure of BKPyV and other polyomaviruses is an intriguing example of this. Through the interaction of host cell transcription and translation machinery with the BKPyV genome and the simple interactions of the capsid proteins, a complex icosahedral-shaped BKPyV virion forms. These simple interactions are essential to the life cycle of PKPyV and our lab found that the use of ODEs made it difficult to model these interactions as well as capture the spatial and temporal heterogeneity of interacting molecules.

As a result, we decided to utilize an alternate modeling technique--agent-based modeling. Agent-based modeling is well suited for modeling emergent properties like viral capsid formation and other molecular interactions. The ABM consists of autonomous agents that represent individual genes, RNAs, and proteins interacting stochastically with each other and their environment based on a set of rules that mimic the biology.

Recently, ABMs have been applied to host pathogen modeling. Duca et al (Duca et al., 2007) created an ABM to produce a virtual model of the tonsils of the nasopharyngeal cavity and peripheral circulation. The host immune response in granuloma formation in response to tuberculosis infection was also modeled using an ABM (Segovia-Juarez et al., 2004). Both these models showed the importance of modeling spatial and temporal dynamics of interacting pathogens and cells.

Host cells essentially determine the growth rate of a virus, and we present an ABM that mimics the salivary gland cell’s support and hindrance of the BKPyV life cycle. We observed that (a) the Boids algorithm can be successfully used for modeling molecular binding, (b) high concentrations of capsid subunits are necessary for virion formation, (c) only small amounts of Tag are necessary for viral replication and (d) there may be salivary gland specific interaction with the regulatory region that enables early region transcription. The model’s source code is available from the GitHub repository, https://github.com/drsuuzzz/assembly.

## Results

Using the Repast Simphony agent-based modeling platform, a single-cell salivary gland model initially infected with one BKPyV virion was designed to investigate the BKPyV replication and assembly process, https://repast.github.io. At present, only known cellular molecules, which affect viral transcription and translation, are represented as agents in the model. Small molecules like calcium, as well as other parameters such as pH, temperature and salt concentrations are currently excluded from the model (Imperiale and Major, 2007; Jiang et al., 2009).

Our model is a simplified representation of a cell, consisting only of nucleus and cytoplasm/endoplasmic reticulum (CER) compartments. There is evidence that polyomavirus capsid proteins and viral DNA co-localize in promyelocytic leukemia nuclear bodies (PML) for viral assembly to occur (Erickson et al., 2012). For the purposes of this model, the nucleus compartment is a simplified representation of a PML domain of the nucleus.

At the start of a simulation, the nuclear compartment contains randomly placed agents representing the BKPyV genome, host Transcription Factors and host DNA Polymerase promoter sites. The CER compartment contains agents representing Ribosomes placed in random locations. Agents will move stochastically within the compartments and may have chance encounters with other agents. These chance encounters may result in the agents binding with one another and the formation of new agents simulating the transcription, translation and replication and assembly of the BKPyV virus.

### Modeling Molecular Binding as a Flock of Birds

Movement and binding of agents is essentially what drives the model and allows the progression of the BKPyV life cycle as most rules will not execute unless an agent is bound to another. For example, a host transcription factor agent binding with the viral genome agent allows the transcription rule to be executed producing an mRNA agent. Biologically, intramolecular interactions are governed by binding affinity/repulsion, stoichiometry, conformational change and biochemical reactions. To simulate this, agents move stochastically, simulating Brownian motion, about the compartment and encounter other agents purely by change. Molecular binding and interactions occur based on these chance encounters and is simulated by the Boids algorithm (Reynolds, 1987), which is summarized here and detailed further in Materials and Methods.

It is challenging, algorithmically, to model independent entities moving together without crowding or overlapping one another as in bird flocking behavior. Boids is a solution to this problem initially proposed and utilized in the computer graphics and animation field. Notwithstanding, bird flocking is somewhat analogous to molecular binding in that bound molecules maintain a distance from each other while moving together without colliding. As such, the Boids implementation in this model consists of simple position calculations and adjustments similar to biochemical electrostatic properties of attraction, repulsion and momentum. In Boids terminology, this is analogous to cohesion, separation and alignment as shown in Figure 2. Repulsion: if the agents are too close together, the agent position is adjusted away from the neighboring agent. Attraction: if the agents are too far apart, the agent position is adjusted towards the neighboring agent. Momentum: the agent position is adjusted towards the calculated average velocity of all neighboring agents. All three calculations together help maintain a separation distance between agents such that they all move together, in the same direction, at the same rate without colliding.

**Figure 2.**
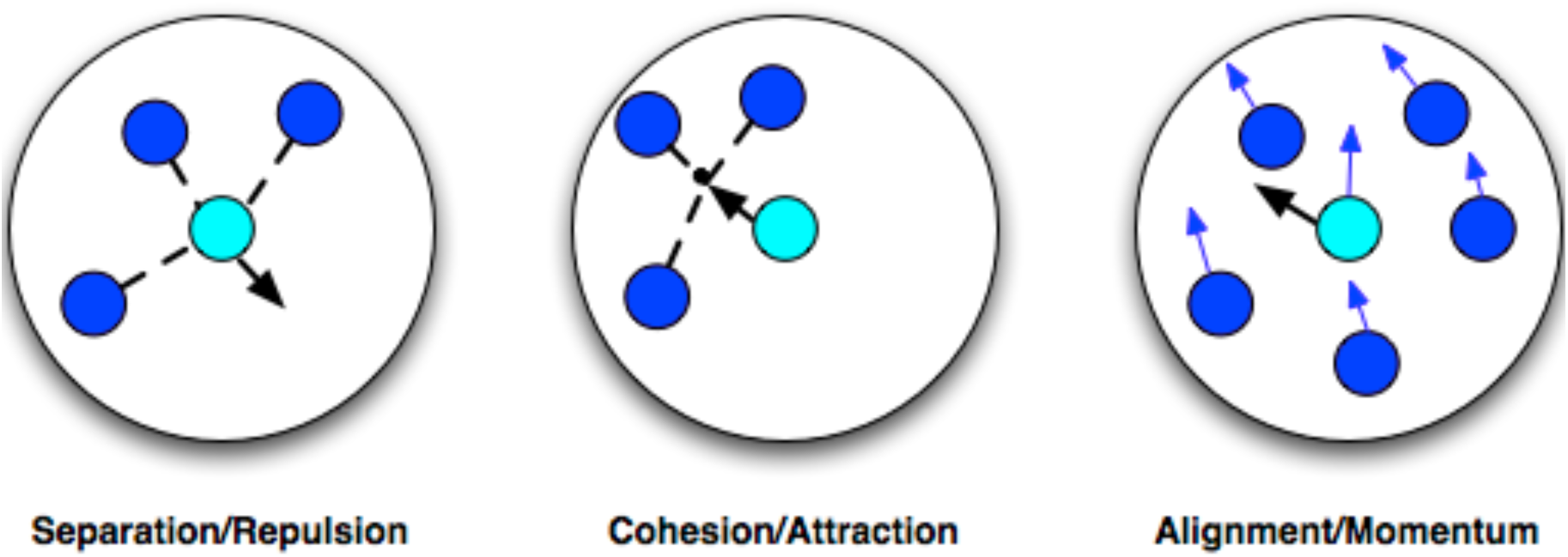
Boids rules. Separation/Repulsion: move away from neighbors. Cohesion/Attraction: move towards average position of neighbors. Alignment/Momentum: move towards average heading of neighbors.

The execution of these rules by the individual agents results in the binding of mRNA agents to ribosome agents to simulate translation, transcription factor agents to promoter agents to simulate transcription and the binding of capsomere agents to form the viral capsid to name a few examples from the ABM.

The most important emergent property resulting from the execution of these rules is the final step in the replication of the BKPyV virus—formation of the viral, icosahedral-shaped capsid-like structure of encapsidated and unencapsidated BKPyV particles. From simple interactions, complex patterns emerge.

### Host Cell Transcription and Translation of BKPyV Proteins

Replication of BKPyV is guided by regulation of the viral genome’s regulatory region, Figure 3a. Transcription follows a somewhat orderly fashion in that first the early region is transcribed which leads to proteins that aid in the replication of the viral genome and finally the transcription of the late region occurs leading to genome encapsidation, Figure 3b. Since host cell machinery is integral to successful viral replication, the model implements host processes in order to explain the differences in replication rates observed between salivary gland and kidney cells.

**Figure 3.**
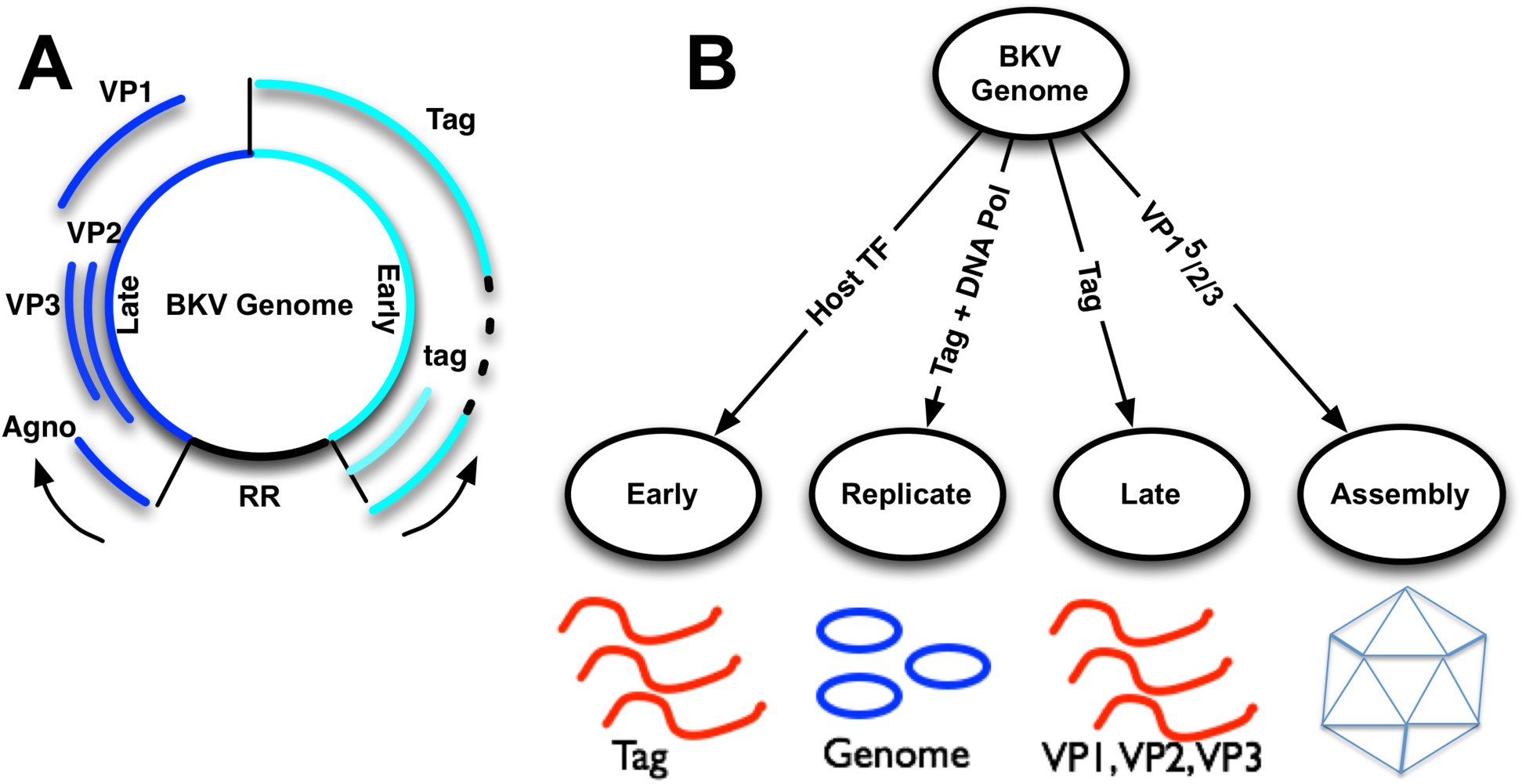
A) BKPyV circular DNA genome depicting the regulatory (RR), early and late regions and transcripts produced. The early region transcribes alternatively spliced RNA Tag or tag mRNA. The late region transcribes alternatively spliced RNAs for translating to VP1, VP2, VP3 and the agnoprotein. B) The model imitates the transcription of early and late regions based on binding to the regulatory region of host transcription factors, Tag or DNA Polymerase. Capsid assembly begins when a VP1 pentamer is bound.

#### Early Transcription

To initiate early region transcription, host transcription factors must be recruited to the regulatory region. Many host transcription factors have been identified which bind to the BKPyV regulatory region and they are represented by a single agent-type in the model (Bethge et al., 2015; Liang et al., 2012; Moens and Vanghelue, 2005). Although we know there is BKPyV tropism for salivary gland cells, it is not known which transcription factors aid early transcription. In our model we set the number of transcription factors to a relatively low number to study rates of virion production. When the transcription factor agent binds with the genome agent, an mRNA agent is created, Figure 4a.

**Figure 4.**
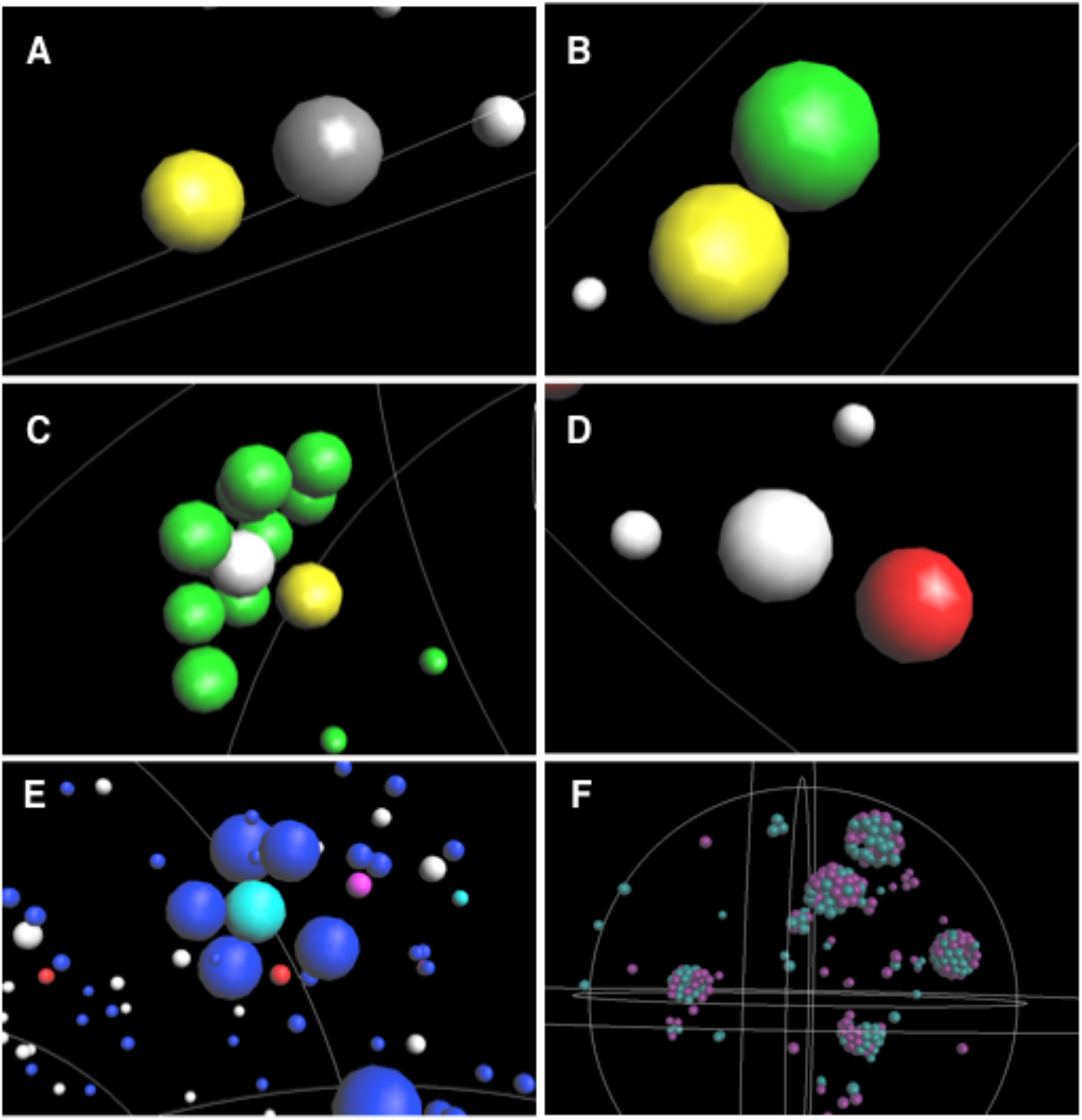
Snapshots of agents in a simulation. A) Early transcription: Host Transcription Factor (Grey) binds with BKPyV Genome (yellow). B) Late transcription: Tag (green) binds with BKPyV Genome (yellow). C) Genome replication: 12 Tag (green) bind with the BKPyV genome (yellow) and recruit DNA Polymerase (light gray). D) Translation: mRNA (red) binds with Ribosome (white). E) Assembly of capsid subunits. 5 VP1 agents (blue) binding to a VP3 agent (cyan). Agents in the background are ribosomes (white), mRNAs (red) and VP2 (magenta). F) A simulation of only capsid self-assembly showing partial capsid formation of VLPs (4) and virions (1). Purple represents VP2 bound VP1 pentamers and green represents VP3 bound VP1 pentamers. The white lines in the simulation indicate the separation of the nucleus and CER components which are represented spherically.

This mRNA agent is an immature mRNA until the alternative splicing rule is executed (Materials and Methods) forming large T antigen (Tag) and small t antigen (tag) transcripts (Eash et al., 2006). Small t antigen (tag) transcripts are ignored at present in the model.

#### Genome Replication

For genome replication, Tag forms a double hexamer and recruits DNA Polymerase to the origin of replication site of the regulatory region (VanLoock et al., n.d.). This is implemented by the accumulation of 6 Tag agents binding to the BKPyV genome agent followed by the binding of a DNA Polymerase agent. A new BKPyV genome agent is eventually created, Figure 3b and Figure 4c. As BKPyV is a DNA virus, the cell must enter S-phase for DNA polymerase to accumulate and initiate BKPyV genome replication (Imperiale and Major, 2007). Tag stimulates entry into S-phase by sequestering pRb resulting in the release of E2F (Lin and Simmons, 1991; Ludlow et al., 1989). In the model, DNA polymerase agents are created by the binding of Tag agents to host DNA fragment agents--an indirect representation of this exact mechanism (Figure 4c).

#### Late Transcription

Finally, late region transcription accompanies BKPyV replication (Imperiale and Major, 2007). This is simulated in the model by the accumulation of Tag agents on the BKPyV genome agent resulting in a late region transcript, Figure 3b and Figure 4b. Alternative splicing is also simulated resulting in complete VP1, VP2 or VP3 transcripts. Agnoprotein is not produced at this initial phase of the model.

#### Translation of viral transcripts

Viral transcripts, mRNA agents, are exported from the nucleus when encountering the boundary between the nucleus and CER compartments. Chance encounters with Ribosome agents with mRNA agents result in the translation of viral protein agents consisting of Tag, VP1, VP2 or VP3, Figure 4d. Tag is imported back into the nucleus when the agent encounters the boundary between the compartments. The model assumes that capsid subunits consisting of VP1 pentamers in complex with either VP2 or VP3 are assembled in the CER before being imported into the nucleus as shown in Figure 4e (Imperiale and Major, 2007). Again, upon encountering the boundary between the two compartments, the capsid subunit complex of a VP1 pentamer and VP2 or VP3 is imported to the nucleus and translated into a single agent for ease of representing the capsid self-assembly process, VP123. The import process described is intended to simulate the nuclear localization signal found on the viral proteins, VP2 or VP3, that is recognized for transfer of the protein complex through the nuclear pore complex (Bennett et al., 2015).

#### Host Transcription Factors affect virion production

The model was tuned to results obtained from BKPyV-salivary gland cell *in vitro* experiments. When monitoring transcript concentrations, a slow ramp-up is observed in the model that coincides with *in vitro* results, Figure 5 (Burger-Calderon et al., 2014; Jeffers-Francis et al., 2015; Jeffers et al., 2009). The correlation of the *in silico* model of VP1 transcript accumulations with *in vitro* data is 0.997. However, Tag transcript and protein correlation with the *in silico* model is −0.84 and 0.48, respectively. In spite of this disagreement, the production of virions has a 0.92 correlation between *in silico* and *in vitro* data. Since the model is producing virions similar to *in vitro* measurements, this implies that the larger quantities of Tag *in vitro* are necessary for host cell interference.

**Figure 5.**
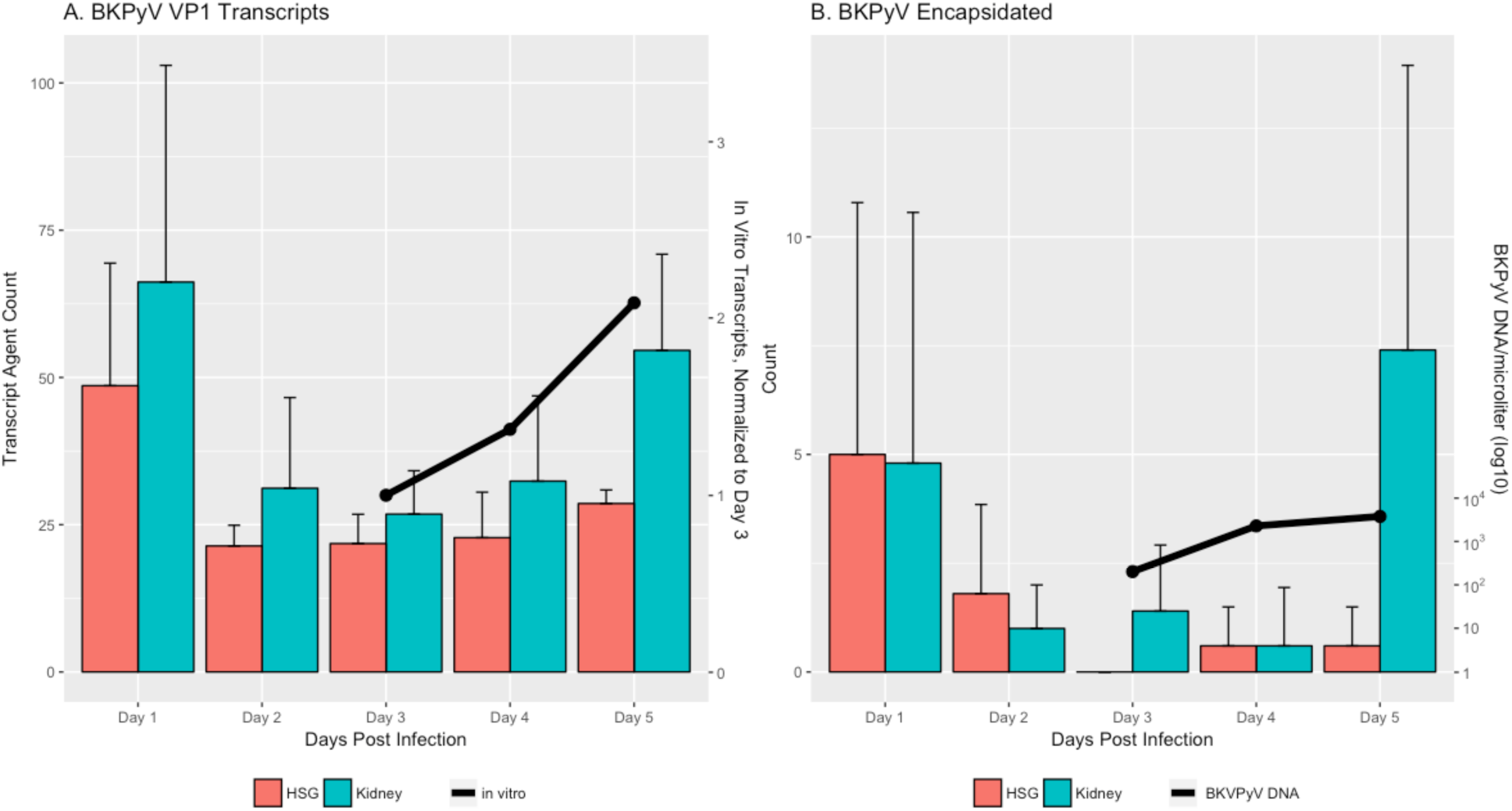
a) VP1 transcripts, b) BKPyV encapsidated agent counts are shown for HSG and Kidney cell simulations. Figures A and B also show in vitro results for comparison.

The replication potential of BKPyV in salivary gland cells has been linked to nucleotide differences in the RR (Burger-Calderon et al., 2014). This suggests that differences in transcription factors expressed between salivary gland cells and kidney cells affect the BKPyV replication efficiency. Although a number of transcription factors that interact with BKPyV have been identified (Bethge et al., 2015; Liang et al., 2012; Moens and Vanghelue, 2005) these studies have been performed in Vero or HRPTEC kidney cell lines. There were a number of approaches that we could of taken to simulate this with the model such as changing probabilities and agent-agent interactions. However, we felt that simply increasing the number of host transcription factor agents more closely represented the biology based on the previously mentioned studies. Figure 5a shows the results from a simulation to mimic the replication rate of a kidney cell where the only parameter changed was the increase in the number of transcription factors from 10 to 20. BKPyV virions were produced within the 24-48 hour window that has been reported in previous *in vitro* studies, Figure 5c. Considering the evidence of salivary gland *in vitro* studies and model simulations, there may exist salivary gland specific mechanisms that enable BKPyV transcription.

To further validate tissue specific expression of transcription factors, 5 normal salivary gland samples from SRP067524 (Bell et al., 2016) and 5 normal kidney samples from SRP060355 (Chhibber et al., 2016) were downloaded and analyzed. Differential expression analysis revealed differences in transcription factor expression mainly in the homeobox family of genes including SIX1 and HOXA7, Supplementary table S1. The Human Protein Atlas also shows differences in gene expression with the same genes between kidney and salivary gland tissues (Fagerberg et al., 2014).

### Self-Assembly of Encapsidated and Unencapsidated BKPyV

The last phase of the replication process is encapsidation of the genome. Self-assembly of the BKPyV virion occurs in the nucleus and is believed to occur more specifically in a PML region (Erickson et al., 2012). The capsid is comprised of the structural proteins VP1, VP2 and VP3 and forms a T=7 icosahedron where 12 VP1 pentamers are located in the 12 vertices of the icosahedron surrounded by 5 neighboring VP1 pentamers (Li et al., 2003; Liddington et al., 1991). 60 pentamers comprise the rest of the structure with 6 neighboring VP1 pentamers for a total of 72 capsomeres (Li et al., 2003; Liddington et al., 1991). A VP1 pentamer is bound with either a VP2 or VP3 protein where the VP2 or VP3 side is presented internally, towards the DNA, and VP1 is on the external surface of the capsid. The C-terminals of the VP1 proteins extend to bind with neighboring VP1 pentamers solidifying the capsid structure. In this model it is assumed that the VP1^5^/VP2 or VP1^5^/VP3 capsomeres are formed in the CER and imported to the nucleus via the VP2 or VP3 nuclear localization signal (NLS). A single agent, VP123, represents the imported capsid subunit.

The model assumes that the capsid subunits can bind DNA and begin assembling around the BKPyV genome agent relying on genome-subunit and subunit-subunit interactions (Roitman-Shemer et al., 2007). However, virus like particles (VLPs) can form in the absence of the viral genome, VP2 and VP3 and thus, aggregation of the subunits leads to the eventual formation of an empty capsid (Li et al., 2003). Empty capsid or VLP formation is modeled by subunit-subunit interactions and an invisible VLP agent that acts as the center of the forming structure in order to enforce the spherical shape of the forming VLP.

As stated previously, a single agent represents a capsid subunit, a VP1 pentamer bound with either VP2 or VP3. Color-coding distinguishes between VP2 or VP3 bound subunits as shown in Figure 4b. Assembly begins upon aggregation of capsid subunits and upon encountering the BKPyV genome agent. Chance encounters of randomly moving capsid agents with the BKPyV genome agent and with other capsid subunit agents allows the gradual formation of the capsid. Once capsid assembly is completed, the structure will continue to move randomly within the nucleus compartment. The spherical, icosahedral structure formed is enforced by the separation distance defined by subunit-subunit and subunit-genome interaction. Egress is currently simulated when the structure eventually encounters the boundary separating the nucleus from the CER compartments. The VLP or virion is then removed from the model and a count of particles is maintained.

The model supports *in vitro* observations of slow accumulations of capsid proteins that eventually lead to BKPyV virions. In addition, our model predicts that a significant accumulation of BKPyV capsomeres is necessary before the BKPyV virions or VLPs are produced (r = 0.91), Figure 6.

**Figure 6.**
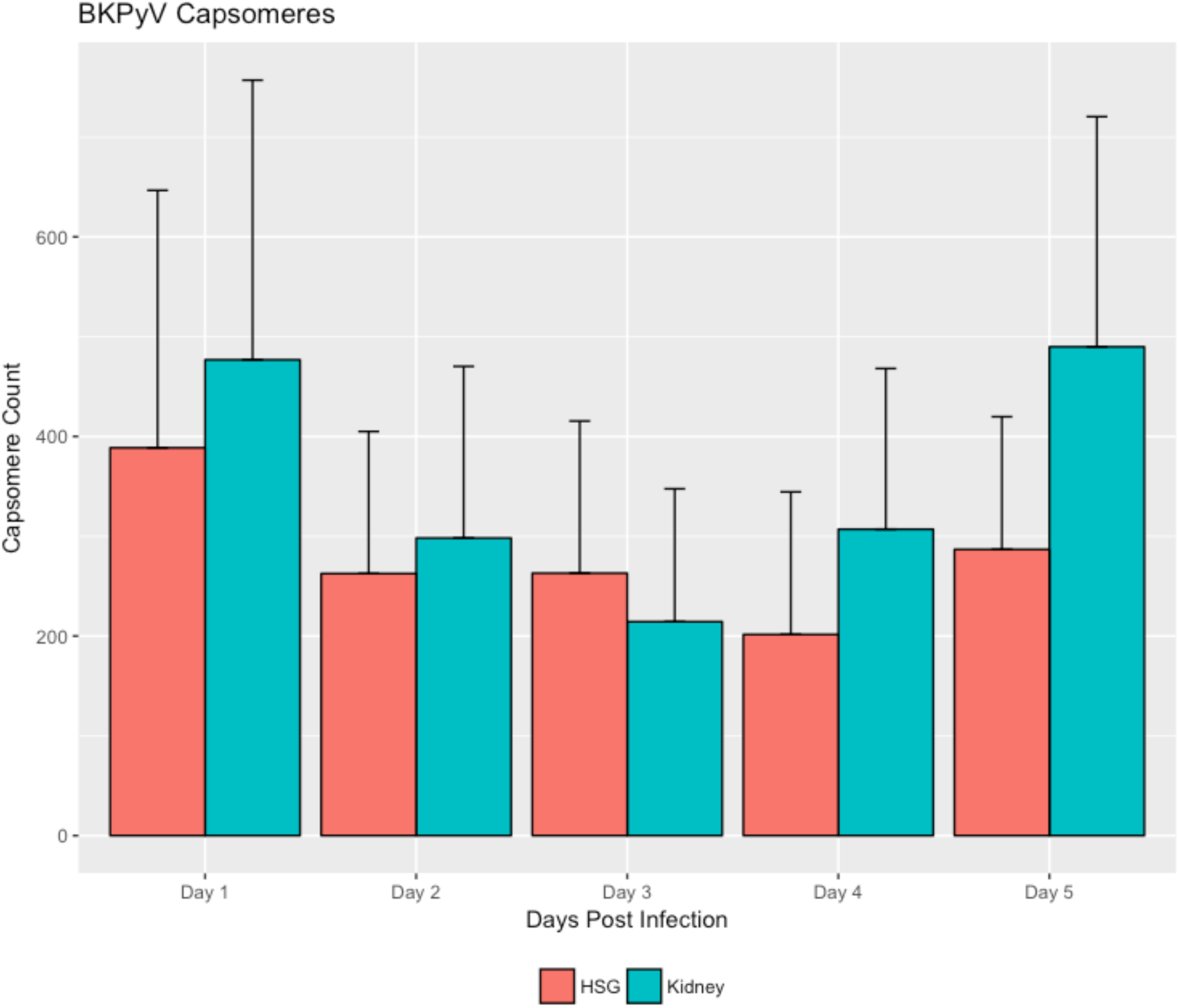
Accumulation of capsomeres (VP1^5^/VP2 or VP1^5^/VP3) correlates with an increase in encapsidated BKPyV virions, r = 0.91.

## Discussion

By modeling molecular interactions within a cell during infection it is possible to understand how the BKPyV affects the function of the cell, to make predictions about therapeutic interventions and to further knowledge about viral pathogenesis in general. While we are aware that there are other factors affecting salivary gland tropism such as entry, uncoating, transport to the nucleus and egress, we have only modeled a portion of the overall viral life cycle—viral replication and assembly. From this initial model we were able to study the viral self-assembly process and the factors that affect successful replication of a BKPyV virion.

The icosahedral structure of BKPyV and other polyomaviruses is intriguing. Although the protein subunits (VP1 pentamers) have a pentagonal shape, they are packed in 6 neighboring subunits (hexavalent) or 5 neighboring subunits (pentavalent) structures (Baker et al., 1989; Griffith et al., 1992; Li et al., 2003; Liddington et al., 1991). Several computational models have been built to understand how nature can form such a complex geometrical structure. One such model is built upon the theory that self-assembly of virus capsids is based on the interaction of structural proteins with neighboring protein subunits. Local rules theory creates a model of the virus capsid based on angles and distances between neighboring subunits approximating the conformation changes of the capsid subunits (Berger et al., 1994; Schwartz et al., 2000, 1998). The mathematical problem of pentagon packing has also been applied to the study of the virus structure making distinctions between loosely packed pentagons as in the case of BKPyV and its polyomavirus cousins and more densely packed pentagons as with papillomaviruses (Tarnai et al., 1995).

Building on this theory, the ABM presented here shows a simple model of viral self-assembly where capsid subunits implement the three Boids rule—separation, cohesion and alignment. The formation of the capsid is an emergent property based on the interaction of capsid subunits with neighboring subunits without the need for angle and distance calculations to force the formation of the viral particle. Transcription and translation, which also rely on the same simple, binding rules, resulted in the production of viral proteins. Further, an additional emergent property of the Boids implementation was a change in the momentum of the forming structure as more and more agents aggregated. Structure movement slowed while moving in random directions as would be observed for larger molecules.

However, for successful viral capsid assembly, it was noted that large accumulation of capsid subunits preceded virion and VLP formation. Muckherjee et al noticed this phenomena when they observed that increased concentrations of capsid subunits encouraged particle formation (Mukherjee et al., 2007). In this model, interacting with the viral genome enforces the T=7 icosahedral curvature of the capsid. Prior work has shown that without the genome, T=1 structures are possible when reducing disulfide bonds and removing calcium ions (Nilsson et al., 2005). These types of structures are possible with this model by simply modifying the separation distances between subunit agents and genome agents.

This model is a proof of principle that can be applied to the study of many other pathogens and cell types as we have done here with salivary gland and kidney cells. Future work for this model consists of implementing the complete BKPyV replication cycle and the addition of therapeutic agents to theorize the affects on viral replication and assembly.

## Materials and Methods

### BKPyV ABM

The ABM was implemented using Repast Simphony version 1.2, https://repast.github.io, a Java based, open-source, ABM framework for developing and visualizing ABMs. All the steps in the life cycle of BKPyV rely on host cell “cooperation”. Thus, a two compartment representation of a host cell with spherical compartments representing the nucleus and CER was designed, Figure 7.

**Figure 7.**
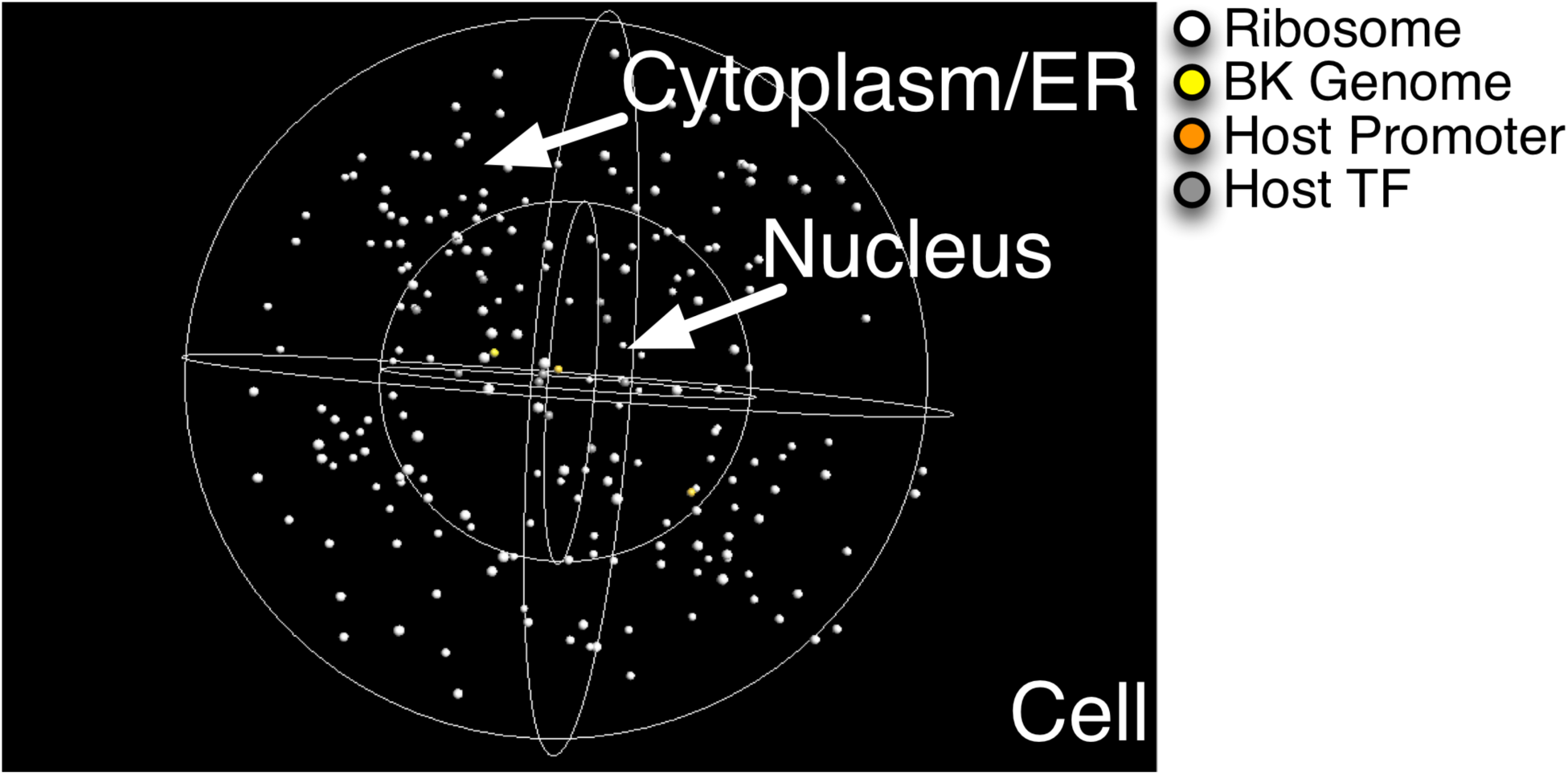
Screen capture of model simulation at initial start.

Each compartment contains agents specific to the compartment as well as agents that can traverse between the two, i.e., mRNA agents exported to the CER from the nucleus, Table 1. The environment of the host cell was represented as a continuous 3-D space such that an agent’s location was represented by its x, y and z-axis floating point coordinates, i.e., agent *b* is located at position (1.1, 2.05, 30.2). When a simulation starts, agents with initial concentrations greater than 1 are placed at random locations within their respective compartments. These initial agents were Ribosome, BK Genome, Host Promoter and Host Transcription Factor (TF), Figure 7 and Table 1.

**Table 1.** Agents, the rules they execute and the agents they interact with

| Agent | Starting Number at t0 | Compartment | Rules | Interacts with |
| --- | --- | --- | --- | --- |
| BK Genome | 1 | Nucleus | move, transcription, egress | Tag, VP123, TF, DNA Pol |
| DNA Pol | 0 | Nucleus | move, death | BK Genome |
| Host Promoter | 2 | Nucleus | move, transcription | Tag |
| mRNA | 0 | CER, Nucleus | move, splice, export, death | Ribosome |
| Ribosome | 280 | CER | move, translation | mRNA |
| Tag | 0 | CER, Nucleus | move, import, death | Host Promoter, BK Genome |
| TF | 10 | Nucleus | move | BK Genome |
| VLP | 0 | Nucleus | move, egress | VP123 |
| VP1 | 0 | CER | move | VP2, VP3 |
| VP123 | 0 | Nucleus | move | BK Genome, VLP |
| VP2 | 0 | CER | move, import | VP1 |
| VP3 | 0 | CER | move, import | VP1 |

In the model, agents represented various host and viral gene products, Table 1. The agents moved based on a simulation of Brownian motion in a 3-dimensional environment described below. Once an agent encountered another agent, interaction rules may be executed, Table 2. In some agent-agent interactions, the movement rule changes to simulate molecular binding by using the Boids algorithm to simulate attraction, repulsion and momentum as detailed further below (Reynolds, 1987).

**Table 2.**
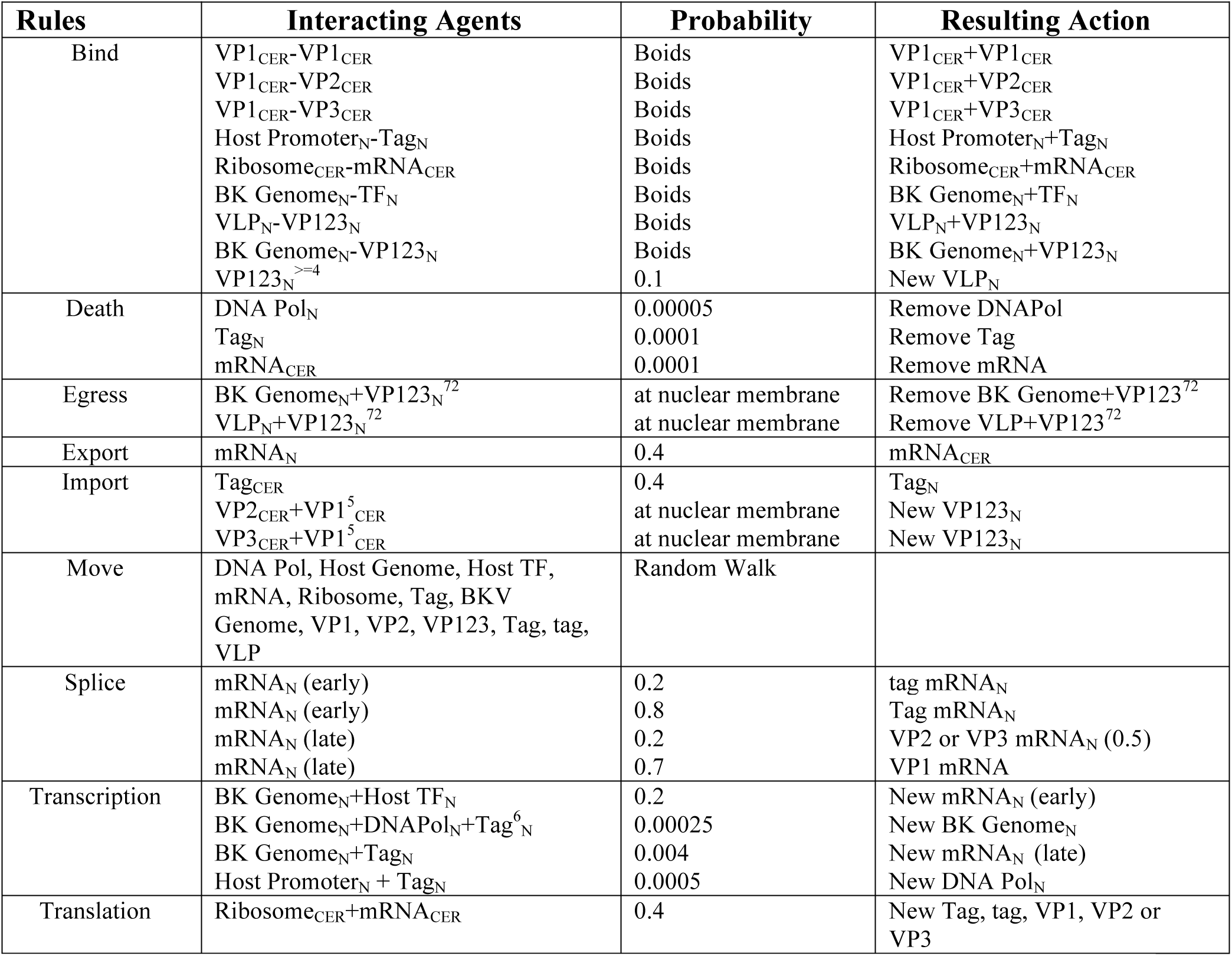
Interacting agents, rule execution probabilities and behavior

### Model Time and Rule Execution

Time in the model was implemented as an iterative cycle where a state update of all agents occurs once per iteration. A time step was completed once all agent rules had been attempted. 60,000 iterations were arbitrarily selected to represent a model day based on repeated simulations during the model parameter estimation and tuning stage.

During a time step, agent rule execution occurred in a random order. A rule was executed once a probability threshold had been met based on a random draw from the uniform distribution as shown in Table 2. For example, the following occurs when the translation rule is executed for a Ribosome agent.

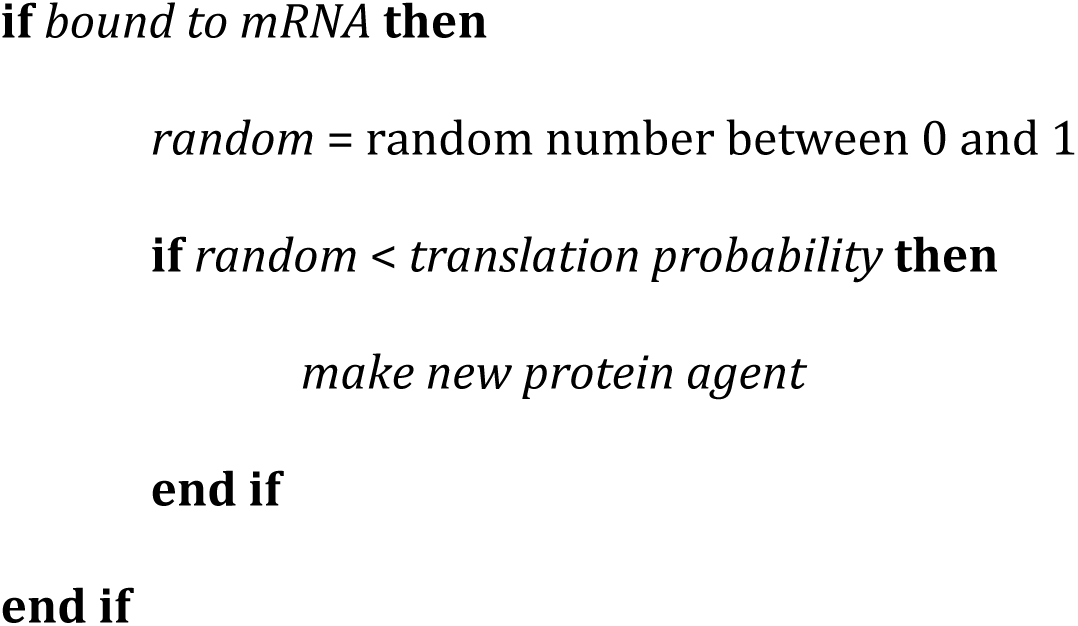

As a result, it could take several time steps before a rule would successfully execute. Probabilities essentially determine how fast or how slow the processes occur within the model.

### Parameter Estimation

There were three important parameter types that were necessary to estimate in this ABM--size of the environment, initial concentration values of agents and rule probabilities. Thus, the parameter space is quite large. Parameters were estimated using a random parameter sweep method, where (1) initial parameter estimates were made based on biological insight, (2) simulations were repeated with randomized parameter values, and (3) parameters were then finalized from simulations that best fit *in vitro* data. As simulations were repeated, rule probabilities were either increased or decreased to speed up or slow down molecular interactions, respectively. Model results were then validated against *in vitro* data by Pearson’s correlation coefficient.

Agent-agent represents unbound neighboring agents, while agent+agent represents bound agents moving together. The agent location is represented as a subscript, i.e., CER or N (nucleus).

### Rules and Agent States

The BKPyV genome is a circular minichromosome, Figure 3a, and can be divided into three regions: early, late and the regulatory regions (RR). The model followed the early and late transcription biological model, Figure 3b, by implementing a simple state machine driven by the BK genome agent and it’s interaction with other agents. The output of a state was either an mRNA or new BK genome agents. The input to a state was an agent binding with the BK genome agent. Successful execution of rules shown in Table 2 resulted in new agents being created (i.e. proteins in the case of translation) or the changing of compartments.

#### Move Rule

All agents implemented the move rule. The move rule is a rather simple simulation of Brownian motion in a 3-dimensional environment where the next position or step the agent makes is determined based on a random draw from a uniform distribution from −0.5 to 0.5 determining the x, y and z coordinates of the direction vector. If a next position calculation would result in an agent overstepping a compartment boundary, it is recalculated to reverse the agent’s direction in order to remain within its compartment. This, in many cases, results in the appearance of the agent "bouncing" off of the compartment boundaries.

#### Transcription Rule

The BK genome agent implemented the transcription rule and its successful execution resulted in the creation of a new early mRNA, late mRNA or new BK genome agent depending on the state of the BK genome agent and agents bound to it. The new agent was then placed within a random location inside a 4 unit radius from the BK genome agent. Once transcription had completed successfully, bound agents would unbind and move freely once again.

#### Splice Rule

The splice rule simulated alternative splicing on an immature early or late mRNA agent. The spliced transcript was determined based on probabilities. If a probability was met, the state of the mRNA was marked complete and its transcript type was set. For early mRNAs, successful execution of the splice rule resulted in the mRNA identifying as either a Tag or tag transcript. For late mRNAs, the splice rule resulted in the mRNA agent identifying as a VP1, VP2 or VP3 transcript.

#### Export Rule

The export rule was only executed for mature (complete) mRNAs when they encountered the boundary between the nucleus and CER containers. Upon successful execution, the agent was moved to the other side of the boundary where it then moved freely within the CER container.

#### Translation Rule

The translation rule was executed after the binding of an mRNA agent to a Ribosome agent. Successful execution of the rule resulted in the creation of a new protein agent depending upon the type of mRNA that was translated. Tag, VP1, VP2 or VP3 agents were created by this rule. tag is not implemented at this time. The new agent was placed within an arbitrarily determined 4 unit radius from the Ribosome agent. The Ribosome agent then unbound from the mRNA agent.

#### Import Rule

The import rule was similar to the export rule in that agents were moved from the CER to the nucleus. The rule was executed when the agent encountered the nucleus boundary.

#### Death Rule

mRNAs are known to randomly degrade, as such a death rule was implemented to mimic this. DNA Pol and Tag also implemented this rule in order to prevent exceedingly high accumulation of the agents and to reduce unnecessary consumption of computational resources.

#### Bind Rule

The bind rule utilized the Boids algorithm to simulate molecular binding (Reynolds, 1987). The simple Boids rules were (i) Separation-steering to avoid crowding, (ii) Alignment-steering towards the average heading of neighbors and (iii) Cohesion-steering to move towards the average position of neighbors. Electrostatically speaking these rules can be interpreted as repulsion (separation) and attraction (cohesion) steps in the binding process of two molecules.

The Boids implementation of the bind rule calculated the next position of an agent based on the position and velocity of neighboring agents. If the agent was too close, the next position was calculated such that it moves away from the other agent a small amount (separation/repulsion). If it is too far away, a position towards the agent was calculated (cohesion/attraction). In addition, the average velocity of the neighboring agents was calculated in order to adjust the velocity of the agent to match its neighbors (alignment/binding). Combined, these rules ensure that the agents moved together in the same direction with the same velocity. However, if an agent made too large of a step when changing position it may have disappeared from a neighbors view and the agent was then unbound.

VP2, VP3 and BK Genome agents only implemented the alignment/binding phase of the Boids algorithm. This was necessary to facilitate binding of neighboring agents on all sides of the agent.

VP1 agents aggregated into pentamers binding with either VP2 or VP3 agents based on successful execution of the Bind rule. Once a VP1^5^/VP2 or VP3 complex was successfully formed and its position was close to the nucleus, successful execution of the import rule caused the removal of the 5 VP1 and associated VP2 or VP3 agents. The VP1^5^/VP2 or VP3 complex was then represented as a single agent, VP123, to facilitate viral self-assembly within the nucleus.

The distance maintained from the genome agent enforced the curvature of the icosahedral shape of the aggregating capsid subunits. To enforce a similar curvature for VLPs, an invisible agent was created when at least 4 capsid subunits began to aggregate. While subunits can bind arbitrarily around the genome agent, subunit-subunit binding was only allowed when forming the VLP around the invisible agent. VP123 agents have a preference for binding with the genome agent over the formation of a VLP.

#### Egress Rule

Lastly, the egress rule was essentially a method to identify whether or not a completed virion or VLP had encountered the nucleus boundary. This rule modeled the as yet unknown process of BKPyV egress from the nucleus. The agents involved in the viral complex were then removed from the model and a count was kept of virions or VLPs produced.

### BKPyV assays

Material and methods for collection of BKPyV data assayed in salivary gland cell lines is as previously described in Jeffers et al (Jeffers-Francis et al., 2015; Jeffers et al., 2009).

### Differential Expression Analysis

Samples were downloaded from the SRA and FASTQs extracted using the SRA Toolkit. FASTQs were quality and adapter trimmed with trim_galore, https://github.com/FelixKrueger/TrimGalore. Alignment and gene quantification of the resulting FASTQs was performed using RSEM (Li and Dewey, 2011) and the hg19 version of the human reference. Expected gene counts were log transformed and upper-quantile normalized. Of the 25 kidney samples, 5 samples were selected based on the principal components analysis and clustering of the upper-quantile normalized gene counts allowing for a balanced statistical analysis with the 5 salivary gland samples. The Bioconductor package sva (Leek and Storey, 2008, 2007) was used to remove any batch effects. Differential expression was determined utilizing a t-test assuming equal variance. The fold change is the difference between group medians.

Transcription factors were identified using information from gene ontology (Ashburner et al., 2000; The Gene Ontology Consortium, 2017), the Unitprot database (The UniProt Consortium, 2017) and the Animal TF DB (Zhang et al., 2015).

## Acknowledgements

A portion of this work was supported by F31DE019594 (SMS) and R21DE023046-01 (JWC) from the National Institute of Dental & Craniofacial Research. The funders had no role in study design, data collection and analysis, decision to publish, or preparation of the manuscript. The content is solely the responsibility of the authors and does not necessarily represent the official views of the National Institute of Dental & Craniofacial Research.

The project was also supported in part by the NIAID UNC Molecular Biology of Viral Diseases Pre-doctoral Training Grant to SMS.

## References

Ashburner, M., Ball, C.A., Blake, J.A., Botstein, D., Butler, H., Cherry, J.M., Davis, A.P., Dolinski, K., Dwight, S.S., Eppig, J.T., Harris, M.A., Hill, D.P., Issel-Tarver, L., Kasarskis, A., Lewis, S., Matese, J.C., Richardson, J.E., Ringwald, M., Rubin, G.M., Sherlock, G., 2000. Gene Ontology: tool for the unification of biology. Nat. Genet. 25, 25.

Baker, T.S., Drak, J., Bina, M., 1989. The capsid of small papova viruses contains 72 pentameric capsomeres: direct evidence from cryo-electron-microscopy of simian virus 40. Biophys. J. 55, 243–253.

Bell, D., Bell, A.H., Bondaruk, J., Hanna, E.Y., Weber, R.S., 2016. In-depth characterization of the salivary adenoid cystic carcinoma transcriptome with emphasis on dominant cell type. Cancer 122, 1513–1522. https://doi.org/10.1002/cncr.29959

Bennett, S.M., Zhao, L., Bosard, C., Imperiale, M.J., 2015. Role of a nuclear localization signal on the minor capsid Proteins VP2 and VP3 in BKPyV nuclear entry. Virology 474, 110–116. https://doi.org/http://dx.doi.org.libproxy.lib.unc.edu/10.1016/j.virol.2014.10.013

Berger, B., Shor, P.W., Tucker-Kellogg, L., King, J., 1994. Local rule-based theory of virus shell assembly. Proc. Natl. Acad. Sci. U. S. A. 91, 7732–7736.

Bethge, T., Hachemi, H.A., Manzetti, J., Gosert, R., Schaffner, W., Hirsch, H.H., 2015. Sp1 Sites in the Noncoding Control Region of BK Polyomavirus Are Key Regulators of Bidirectional Viral Early and Late Gene Expression. J. Virol. 89, 3396–3411. https://doi.org/10.1128/JVI.03625-14

Burger-Calderon, R., Madden, V., Hallett, R.A., Gingerich, A.D., Nickeleit, V., Webster-Cyriaque, J., 2014. Replication of Oral BK Virus in Human Salivary Gland Cells. J. Virol. 88, 559–573. https://doi.org/10.1128/JVI.02777-13

Chhibber, A., French, C.E., Yee, S.W., Gamazon, E.R., Theusch, E., Qin, X., Webb, A., Papp, A.C., Wang, A., Simmons, C.Q., Konkashbaev, A., Chaudhry, A.S., Mitchel, K., Stryke, D., Ferrin, T.E., Weiss, S.T., Kroetz, D.L., Sadee, W., Nickerson, D.A., Krauss, R.M., George, A.L., Schuetz, E.G., Medina, M.W., Cox, N.J., Scherer, S.E., Giacomini, K.M., Brenner, S.E., 2016. Transcriptomic variation of pharmacogenes in multiple human tissues and lymphoblastoid cell lines. Pharmacogenomics J. 17, 137.

Drachenberg, C.B., Papadimitriou, J.C., Wali, R., Cubitt, C.L., Ramos, E., 2003. BK Polyoma Virus Allograft Nephropathy: Ultrastructural Features from Viral Cell Entry to Lysis. Am. J. Transplant. 3, 1383–1392.

Duca, K.A., Shapiro, M., Delgado-Eckert, E., Hadinoto, V., Jarrah, A.S., Laubenbacher, R., Lee, K., Luzuriaga, K., Polys, N.F., Thorley-Lawson, D.A., 2007. A Virtual Look at Epstein-Barr Virus Infection: Biological Interpretations. PLoS Pathog. 3, e137.

Dugan, A.S., Eash, S., Atwood, W.J., 2005. An N-Linked Glycoprotein with {alpha}(2,3)-Linked Sialic Acid Is a Receptor for BK Virus. J. Virol. 79, 14442–14445. https://doi.org/10.1128/JVI.79.22.14442-14445.2005

Eash, S., Atwood, W.J., 2005. Involvement of Cytoskeletal Components in BK Virus Infectious Entry. J. Virol. 79, 11734–11741. https://doi.org/10.1128/JVI.79.18.11734-11741.2005

Eash, S., Manley, K., Gasparovic, M., Querbes, W., Atwood, W., 2006. The human polyomaviruses. Cell. Mol. Life Sci. 63, 865–876.

Eash, S., Querbes, W., Atwood, W.J., 2004. Infection of Vero Cells by BK Virus Is Dependent on Caveolae. J. Virol. 78, 11583–11590. https://doi.org/10.1128/JVI.78.21.11583-11590.2004

Erickson, K.D., Bouchet-Marquis, C., Heiser, K., Szomolanyi-Tsuda, E., Mishra, R., Lamothe, B., Hoenger, A., Garcea, R.L., 2012. Virion Assembly Factories in the Nucleus of Polyomavirus-Infected Cells. PLoS Pathog 8, e1002630.

Fagerberg, L., Hallström, B.M., Oksvold, P., Kampf, C., Djureinovic, D., Odeberg, J., Habuka, M., Tahmasebpoor, S., Danielsson, A., Edlund, K., Asplund, A., Sjöstedt, E., Lundberg, E., Szigyarto, C.A.-K., Skogs, M., Takanen, J.O., Berling, H., Tegel, H., Mulder, J., Nilsson, P., Schwenk, J.M., Lindskog, C., Danielsson, F., Mardinoglu, A., Sivertsson, Å., von Feilitzen, K., Forsberg, M., Zwahlen, M., Olsson, I., Navani, S., Huss, M., Nielsen, J., Ponten, F., Uhlén, M., 2014. Analysis of the Human Tissue-specific Expression by Genome-wide Integration of Transcriptomics and Antibody-based Proteomics. Mol. Cell. Proteomics 13, 397–406. https://doi.org/10.1074/mcp.M113.035600

Gerits, N., Johannessen, M., TÃ¼mmler, C., Walquist, M., Kostenko, S., Snapkov, I., van Loon, B., Ferrari, E., HÃ¼bscher, U., Moens, U., 2015. Agnoprotein of polyomavirus BK interacts with proliferating cell nuclear antigen and inhibits DNA replication. Virol. J. 12, 1–14. https://doi.org/10.1186/s12985-014-0220-1

Griffith, J.P., Griffith, D.L., Rayment, I., Murakami, W.T., 1992. Inside polyomavirus at 25-Aa resolution. Nature 355, 652–654.

Imperiale, M.J., Major, E.O., 2007. Polyomaviruses, in: Fields, B.N., Knipe, D.M., Howley, P.M. (Eds.), Fields’ Virology. Lippincott Williams & Wilkins, Philadelphia, PA, pp. 2263–2298.

Inoue, T., Dosey, A., Herbstman, J.F., Ravindran, M.S., Skiniotis, G., Tsai, B., 2015. ERdj5 Reductase Cooperates with Protein Disulfide Isomerase To Promote Simian Virus 40 Endoplasmic Reticulum Membrane Translocation. J. Virol. 89, 8897–8908.

Jeffers-Francis, L.K., Burger-Calderon, R., Webster-Cyriaque, J., 2015. Effect of Leflunomide, Cidofovir and Ciprofloxacin on replication of BKPyV in a salivary gland in vitro culture system. Antiviral Res. 118, 46–55. https://doi.org/http://dx.doi.org.libproxy.lib.unc.edu/10.1016/j.antiviral.2015.02.002

Jeffers, L.K., Madden, V., Webster-Cyriaque, J., 2009. BK virus has tropism for human salivary gland cells in vitro: Implications for transmission. Virology 394, 183–193. https://doi.org/10.1016/j.virol.2009.07.022

Jiang, M., Abend, J.R., Tsai, B., Imperiale, M.J., 2009. Early events during BK virus entry and disassembly. J. Virol. 83, 1350–1358.

Johannessen, M., Myhre, M., Dragset, M., Tummler, C., Moens, U., 2008. Phosphorylation of human polyomavirus BK agnoprotein at Ser-11 is mediated by PKC and has an important regulative function. Virology 379, 97–109.

Kepler, G.M., Nguyen, H.K., Webster-Cyriaque, J., Banks, H.T., 2007. A dynamic model for induced reactivation of latent virus. J. Theor. Biol. 244, 451–462.

Leek, J.T., Storey, J.D., 2008. A general framework for multiple testing dependence. Proc. Natl. Acad. Sci. 105, 18718 LP–18723. https://doi.org/10.1073/pnas.0808709105

Leek, J.T., Storey, J.D., 2007. Capturing Heterogeneity in Gene Expression Studies by Surrogate Variable Analysis. PLOS Genet. 3, e161.

Li, B., Dewey, C., 2011. RSEM: accurate transcript quantification from RNA-Seq data with or without a reference genome. BMC Bioinformatics 12, 323.

Li, T.-C., Takeda, N., Kato, K., Nilsson, J., Xing, L., Haag, L., Cheng, R.H., Miyamura, T., 2003. Characterization of self-assembled virus-like particles of human polyomavirus BK generated by recombinant baculoviruses. Virology 311, 115–124.

Liang, B., Tikhanovich, I., Nasheuer, H.P., Folk, W.R., 2012. Stimulation of BK Virus DNA Replication by NFI Family Transcription Factors. J. Virol. 86, 3264–3275. https://doi.org/10.1128/JVI.06369-11

Liddington, R.C., Yan, Y., Moulai, J., Sahli, R., Benjamin, T.L., Harrison, S.C., 1991. Structure of simian virus 40 at 3.8-A resolution. Nature 354, 278–284.

Lin, J.Y., Simmons, D.T., 1991. The ability of large T antigen to complex with p53 is necessary for the increased life span and partial transformation of human cells by simian virus 40. J. Virol. 65, 6447–6453.

Low, J.A., Magnuson, B., Tsai, B., Imperiale, M.J., 2006. Identification of Gangliosides GD1b and GT1b as Receptors for BK Virus. J. Virol. 80, 1361–1366. https://doi.org/10.1128/JVI.80.3.1361-1366.2006

Ludlow, J.W., DeCaprio, J.A., Huang, C.-M., Lee, W.-H., Paucha, E., Livingston, D.M., 1989. SV40 large T antigen binds preferentially to an underphosphorylated member of the retinoblastoma susceptibility gene product family. Cell, 56, 57–65.

Moens, U., Vanghelue, M., 2005. Polymorphism in the genome of non-passaged human polyomavirus BK: implications for cell tropism and the pathological role of the virus. Virology 331, 209–231.

Moriyama, T., Marquez, J.P., Wakatsuki, T., Sorokin, A., 2007. Caveolar Endocytosis Is Critical for BK Virus Infection of Human Renal Proximal Tubular Epithelial Cells. J. Virol. 81, 8552–8562. https://doi.org/10.1128/JVI.00924-07

Moriyama, T., Sorokin, A., 2008. Intracellular trafficking pathway of BK Virus in human renal proximal tubular epithelial cells. Virology, 371, 336–349.

Mukherjee, S., Abd-El-Latif, M., Bronstein, M., Ben-nun-Shaul, O., Kler, S., Oppenheim, A., 2007. High cooperativity of the SV40 major capsid protein VP1 in virus assembly. PLoS One 2, e765.

Nickeleit, V., Singh, H.K., Mihatsch, M.J., 2003. Polyomavirus nephropathy: morphology, pathophysiology, and clinical management. Curr. Opin. Nephrol. Hypertens. 12, 599–605.

Nilsson, J., Miyazaki, N., Xing, L., Wu, B., Hammar, L., Li, T.C., Takeda, N., Miyamura, T., Cheng, R.H., 2005. Structure and assembly of a T=1 virus-like particle in BK polyomavirus. J. Virol. 79, 5337–5345.

Okada, Y., Suzuki, T., Sunden, Y., Orba, Y., Kose, S., Imamoto, N., Takahashi, H., Tanaka, S., Hall, W.W., Nagashima, K., Sawa, H., 2005. Dissociation of heterochromatin protein 1 from lamin B receptor induced by human polyomavirus agnoprotein: role in nuclear egress of viral particles. EMBO Rep. 6, 452–457.

Padgett, B.L., Walker, D.L., 1976. New human papovaviruses. Prog. Med. Virol. 22, 1–35.

Pastrana, D. V, Ray, U., Magaldi, T.G., Schowalter, R.M., Çuburu, N., Buck, C.B., 2013. BK

Polyomavirus Genotypes Represent Distinct Serotypes with Distinct Entry Tropism. J. Virol. 87, 10105–10113. https://doi.org/10.1128/JVI.01189-13

Perelson, A.S., Neumann, A.U., Markowitz, M., Leonard, J.M., Ho, D.D., 1996. HIV-1 Dynamics in Vivo: Virion Clearance Rate, Infected Cell Life-Span, and Viral Generation Time. Science (80-.). 271, 1582–1586. https://doi.org/10.1126/science.271.5255.1582

Reynolds, C.W., 1987. Flocks, Herds, and Schools: A Distributed Behavioral Model. Comput. Graph. (ACM). 21, 25–34.

Ribeiro, R.M., Lo, A., Perelson, A.S., 2002. Dynamics of hepatitis B virus infection. Microbes Infect. 4, 829–835.

Roitman-Shemer, V., Stokrova, J., Forstova, J., Oppenheim, A., 2007. Assemblages of simian virus 40 capsid proteins and viral DNA visualized by electron microscopy. Biochem. Biophys. Res. Commun. 353, 424–430.

Schwartz, R., Garcea, R.L., Berger, B., 2000. “Local Rules□7 Theory Applied to Polyomavirus Polymorphic Capsid Assemblies. Virology 268, 461–470.

Schwartz, R., Shor, P.W., Prevelige, P.E., Berger, B., 1998. Local rules simulation of the kinetics of virus capsid self-assembly. Biophys. J. 75, 2626–2636.

Segovia-Juarez, J.L., Ganguli, S., Kirschner, D., 2004. Identifying control mechanisms of granuloma formation during M. tuberculosis infection using an agent-based model. J. Theor. Biol. 231, 357–376.

Suzuki, T., Orba, Y., Okada, Y., Sunden, Y., Kimura, T., Tanaka, S., Nagashima, K., Hall, W.W., Sawa, H., 2010. The Human Polyoma JC Virus Agnoprotein Acts as a Viroporin. PLoS Pathog 6, e1000801.

Tarnai, T., Gaspar, Z., Szalai, L., 1995. Pentagon packing models for “all-pentamer” virus structures. Biophys. J. 69, 612–618.

The Gene Ontology Consortium, 2017. Expansion of the Gene Ontology knowledgebase and resources. Nucleic Acids Res. 45, D331–D338.

The UniProt Consortium, 2017. UniProt: the universal protein knowledgebase. Nucleic Acids Res. 45, D158–D169.

Tognon, M., Corallini, A., Martini, F., Negrini, M., Barbanti-Brodano, G., 2003. Oncogenic transformation by BK virus and association with human tumors. Oncogene 22, 5192–5200.

VanLoock, M.S., Alexandrov, A., Yu, X., Cozzarelli, N.R., Egelman, E.H., n.d. SV40 Large T Antigen Hexamer Structure. Curr. Biol. 12, 472–476. https://doi.org/10.1016/S0960-9822(02)00696-6

Zhang, H.-M., Liu, T., Liu, C.-J., Song, S., Zhang, X., Liu, W., Jia, H., Xue, Y., Guo, A.-Y., 2015. AnimalTFDB 2.0: a resource for expression, prediction and functional study of animal transcription factors. Nucleic Acids Res. 43, D76–D81.

